# Efficient variant set mixed model association tests for continuous and binary traits in large-scale whole genome sequencing studies

**DOI:** 10.1101/395046

**Authors:** Han Chen, Jennifer E. Huffman, Jennifer A. Brody, Chaolong Wang, Seunggeun Lee, Zilin Li, Stephanie M. Gogarten, Tamar Sofer, Lawrence F. Bielak, Joshua C. Bis, John Blangero, Russell P. Bowler, Brian E. Cade, Michael H. Cho, Adolfo Correa, Joanne E. Curran, Paul S. de Vries, David C. Glahn, Xiuqing Guo, Andrew D. Johnson, Sharon Kardia, Charles Kooperberg, Joshua P. Lewis, Xiaoming Liu, Rasika A. Mathias, Braxton D. Mitchell, Jeffrey R. O’Connell, Patricia A. Peyser, Wendy S. Post, Alex P. Reiner, Stephen S. Rich, Jerome I. Rotter, Edwin K. Silverman, Jennifer A. Smith, Ramachandran S. Vasan, James G. Wilson, Lisa R. Yanek, NHLBI Trans-Omics for Precision Medicine (TOPMed) Consortium, TOPMed Hematology and Hemostasis Working Group, Susan Redline, Nicholas L. Smith, Eric Boerwinkle, Ingrid B. Borecki, L. Adrienne Cupples, Cathy C. Laurie, Alanna C. Morrison, Kenneth M. Rice, Xihong Lin

## Abstract

With advances in Whole Genome Sequencing (WGS) technology, more advanced statistical methods for testing genetic association with rare variants are being developed. Methods in which variants are grouped for analysis are also known as variant-set, gene-based, and aggregate unit tests. The burden test and Sequence Kernel Association Test (SKAT) are two widely used variant-set tests, which were originally developed for samples of unrelated individuals and later have been extended to family data with known pedigree structures. However, computationally-efficient and powerful variant-set tests are needed to make analyses tractable in large-scale WGS studies with complex study samples. In this paper, we propose the variant-Set Mixed Model Association Tests (SMMAT) for continuous and binary traits using the generalized linear mixed model framework. These tests can be applied to large-scale WGS studies involving samples with population structure and relatedness, such as in the National Heart, Lung, and Blood Institute’s Trans-Omics for Precision Medicine (TOPMed) program. SMMAT tests share the same null model for different variant sets, and a virtue of this null model, which includes covariates only, is that it needs to be only fit once for all tests in each genome-wide analysis. Simulation studies show that all the proposed SMMAT tests correctly control type I error rates for both continuous and binary traits in the presence of population structure and relatedness. We also illustrate our tests in a real data example of analysis of plasma fibrinogen levels in the TOPMed program (n = 23,763), using the Analysis Commons, a cloud-based computing platform.

## INTRODUCTION

In recent years, massive DNA sequence data have been generated. Large-scale whole genome sequencing projects, such as the National Heart, Lung, and Blood Institute’s (NHLBI) Trans-Omics for Precision Medicine (TOPMed) program and the National Human Genome Research Institute’s (NHGRI) Genome Sequencing Project (GSP), have produced whole genome sequences from over 120,000 samples. The designs of the studies from which participants are drawn need not be uniform or simple; for example, TOPMed includes population-based cohorts, family studies, and case-control studies, some of which are conducted in recently admixed populations, and some of which involve large pedigrees of closely-related participants.

In population-based cohorts and case-control studies, population stratification and cryptic relatedness are major sources of confounding that need to be accounted for in association tests. For common single variant analysis, linear mixed models that use an estimated genetic relationship matrix (GRM) to account for both population stratification and cryptic relatedness have been widely applied in Genome-Wide Association Studies (GWAS) to analyze structured and related samples.^1-6^ For binary traits, however, we previously showed that linear mixed models may not be appropriate in the presence of population stratification due to misspecified mean-variance relationships. Therefore, we instead proposed a computationally efficient method GMMAT^7^ to perform single common variant tests in GWAS by fitting generalized linear mixed models (GLMMs),^8^ which simultaneously account for population structure, cryptic relatedness, and shared environmental effects, using multiple variance components and/or random effects.

Hundreds of millions of genetic variants, mostly with a low and extremely rare minor allele frequency (MAF), are being analyzed in large-scale sequencing projects such as TOPMed and GSP. Yet, single-variant tests that have been widely used in GWAS are generally underpowered for analyzing rare genetic variants from sequencing studies. To circumvent this problem, statistical tests such as the burden test,^9-12^ Sequence Kernel Association Test (SKAT),^13^ and their various combinations^14-16^ have been proposed. These tests analyze multiple genetic variants in sets, grouped by genes, genomic regions, or other bioinformatic aggregation units. Most of these tests were originally developed to analyze samples from unrelated individuals, as well as extensions to analyze family data with known pedigree structures in the parametric mixed model and semiparametric generalized estimating equation frameworks.^17-23^ However, these existing methods do not account for cryptic relatedness and have not been applied to large-scale whole genome sequencing studies with population structure, familial and/or cryptic relatedness, due to statistical and computational challenges.

One challenge is that among traditional variant set tests such as burden tests and SKAT, no single approach is uniformly most powerful. Another challenge is that existing hybrid tests that combine burden tests and SKAT, such as SKAT-O,^14^ MiST^15^ and aSPU,^16^ are powerful but are subject to much greater computational loads than either the burden test or SKAT alone in the GLMM framework. Of note, SKAT-O is slower than SKAT because it searches on a grid for the optimal linear combination of the burden test and SKAT statistics. MiST requires adjusting for the genetic burden as a covariate in the SKAT model, and hence needs to fit a burden model for each variant set. In large samples of possibly related individuals, extension of MiST is not as practical as in unrelated samples, since fitting a mixed effects model using the burden score for each variant set (or each test unit) is computationally intensive across the genome. Finally, aSPU uses a permutation or Monte Carlo simulation procedure to compute the p values, which can also be challenging in the context of large-scale whole genome sequencing studies with both population structure and relatedness. Therefore, there is a pressing need to develop powerful and computationally-efficient statistical methods for large-scale whole genome sequencing studies.

To address these statistical and computational challenges, we develop the variant Set Mixed Model Association Tests (SMMAT), computationally-efficient variant set tests for both continuous and binary traits, which are applicable to large-scale whole genome sequencing studies with structured and related samples. We include four tests in the SMMAT framework: the burden test (SMMAT-B), SKAT (SMMAT-S), SKAT-O (SMMAT-O), and an efficient hybrid test to combine the burden test and SKAT (SMMAT-E). All the four SMMAT tests share the same reduced model under the null hypothesis, i.e., the GLMM with only covariates, which only needs to be fit once for all genetic variant sets in an analysis. We show that all of these tests can be constructed using shared single-variant scores and their covariance matrices, thus further improving the computational efficiency in practice compared to performing these tests separately. Moreover, it has been shown that single-variant scores and their covariance matrices can also be used in the meta-analysis of variant set tests,^24, 25^ thus SMMAT can be directly applied to combine multi-cohort studies ranging from unstructured independent samples, to structured and related samples. Finally, we develop a unified analysis pipeline in our software package Generalized linear Mixed Model Association Tests (GMMAT) that implements SMMAT variant set tests in both single study (pooled analysis) and meta-analysis contexts to facilitate research on rare genetic variants from large-scale sequencing studies. We demonstrate the application of our method to the analysis of fibrinogen levels in the TOPMed study.

## METHODS

### Generalized Linear Mixed Models (GLMMs)

We formulate the SMMAT tests (SMMAT-B, SMMAT-S, SMMAT-O and SMMAT-E) from the same GLMM

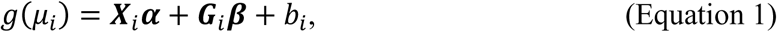

where *g*(·) is a monotonic “link” function that connects the mean of phenotype *y*_*i*_, denoted by *μ*_*i*_ = *E*(*y*_*i*_|***X***_*i*_, ***G***_*i*_, *b*_*i*_), for subject *i* of *n* samples, to the covariate row vector ***X***_*i*_, the genotype row vector ***G***_*i*_ for *q* genetic variants in a set, and the random effects *b*_*i*_ that accounts for population structure and relatedness. The phenotypes *y*_*i*_ follow a distribution in the exponential family. For continuous traits, we usually assume *y*_*i*_ follow a normal distribution and use an identity link function; for binary traits, we assume *y*_*i*_ follow a Bernoulli distribution and use a logit link function. In Equation 1, ***α*** is a *p* × 1 vector of fixed covariate effects including an intercept, and the genotype effects ***β*** are assumed to be a *q* × 1 vector whose distribution has mean ***W*1**_*q*_*β*_0_ and covariance *θ****W***^2^, where ***W*** = *diag*{*w*_*j*_} is a pre-specified *q* × *q* matrix assigning weights to each variant, *θ* is a variance component parameter, and **1**_*q*_ is a column vector of length *q* with all elements 1. We assume that 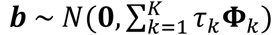 is an *n* × 1 vector of random effects *b*_*i*_, with variance component parameters *τ*_*k*_ and known *n* × *n* relatedness matrices **Φ**_*k*_. We allow for multiple random effects to account for complex sampling designs such as hierarchical designs and shared environmental effects.

### SMMAT-B, SMMAT-S and SMMAT-O

In Equation 1, testing the genotype effects of *q* variants *H*_0_: ***β*** = **0** is equivalent to testing the null hypothesis that *H*_0_: *β*_0_ = 0 and *θ* = 0. The reduced GLMM under this null hypothesis specifies that

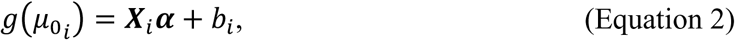

where 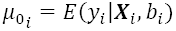. If we test *H*_0_: *β*_0_ = 0 under the assumption that *θ* = 0, a burden score test SMMAT-B can be constructed as

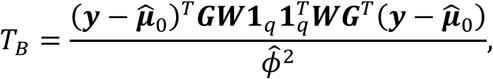

where ***y*** = (*y*_1_ *y*_2_ … *y*_*n*_)^*T*^ is an *n* × 1 vector of phenotypes *y*_*i*_, 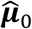 is a vector of fitted mean values under the model in Equation 2, 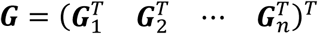 is an *n* × *q* genotype matrix of the variant set in the test, and 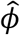 is an estimate of the dispersion parameter (or the residual variance) *Φ*. Under *H*_0_: *β*_0_ = 0, the statistic *T*_*B*_ asymptotically follows 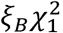, where the scalar 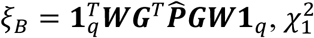 is a chi-square distribution with 1 df, and 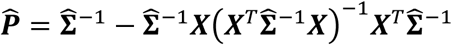 is the *n* × *n* projection matrix of the null GLMM (Equation 2), 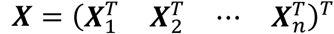 is an *n* × *p* covariate matrix, 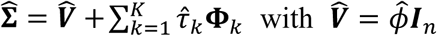 for continuous traits in linear mixed models, and 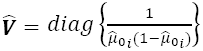 for binary traits in logistic mixed models (where the dispersion parameter *Φ* is known to be 1).

On the other hand, if we test *H*_0_: *θ* = 0 under the assumption *β*_0_ = 0, a variance component score-type test SMMAT-S can be constructed as

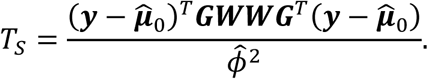

Under *H*_0_: *θ* = 0, *T*_*S*_ asymptotically follows 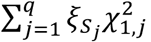, where 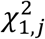 are independent chi-square distributions with 1 df, and 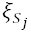 are the eigenvalues of 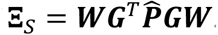.

If one assumes *β*_0_ has mean 0 and variance *γ*, ***β*** then follows a distribution 0 and covariance 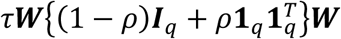, where *τ* = *γ* + *θ* and *ρ* = *Σ*/(*Σ* + *θ*), which takes values between 0 and 1. The joint null hypothesis *H*_0_: *β*_0_ = 0 and *θ* = 0 is equivalent to *H*_0_: *τ* = 0. Given *ρ*, a variance component score-type test can be constructed as

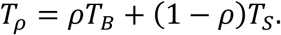

If *ρ* = 1, *T*_*ρ*_ becomes the SMMAT-B burden statistic *T*_*B*_, which assumes ***β*** are the same for all *q* variants after weighting. If *ρ* = 0, T_ρ_ becomes the SMMAT-S SKAT statistic *T*_*S*_. If an optimal *ρ* is obtained by minimizing the p-value of *T*_*ρ*_, then SMMAT-O can be constructed, with its p value calculated using a one-dimensional numerical integration, following SKAT-O.^14^ A key advantage of SMMAT-O is that it maximizes the power by using the optimal linear combination of the mixed model burden test SMMAT-B and the mixed model SKAT SMMAT-S. As it requires a grid search over *ρ*, it is computationally considerably more expensive than SMMAT-B and SMMAT-S. We propose in the next section a computationally much more efficient method to combine SMMAT-B and SMMAT-S.

### SMMAT-E

An alternative joint test to SMMAT-O for *H*_0_: *β*_0_ = 0 and *θ* = 0 can be constructed using two asymptotically independent tests: a test for *H*_0_: *β*_0_ = 0 versus *H*_1_: *β*_0_ ≠ 0 under the constraint *θ* = 0, and a test for *H*_0_: *θ* = 0 versus *H*_1_: *θ* > 0 with *β*_0_ as a nuisance parameter that is estimated under *H*_0_: *θ* = 0. In unrelated samples, this testing strategy is MiST,^15^ which requires the burden model to be fit for each SNP set. We note that the first test is SMMAT-B *T*_*B*_ in the SMMAT framework, and the second test *T*_*θ*_ can be constructed from the null burden GLMM

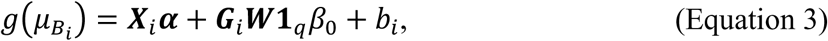

where 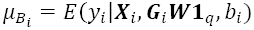 is the mean of *y*_*i*_ in the burden GLMM. We can construct a SKAT-type statistic adjusting for the genetic burden

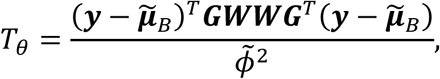

where 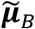 is a vector of fitted values 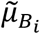 using the burden GLMM in Equation 3 for a given variant set. However, fitting this burden GLMM separately for each variant set is computationally expensive in large-scale whole-genome association studies.

Therefore, we propose a different computationally efficient strategy by assuming that the mean of genetic effects *β*_0_ is not large, a reasonable assumption for most genomic regions and most complex human diseases. Then we can construct *T*_*θ*_ efficiently without refitting the burden GLMMs in Equation 3 for each variant set across the genome. We show in the Appendix that *T*_*θ*_ can be approximated by

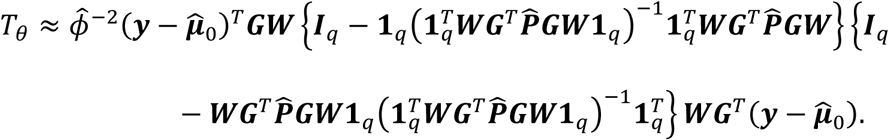

Therefore, under *H*_0_: *θ* = 0, *T*_*θ*_ asymptotically approximately follows 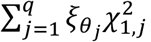, where 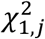 are independent chi-square distributions with 1 df, and 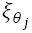 are the eigenvalues of 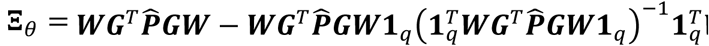. By the central limit theorem, both 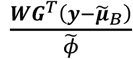 and 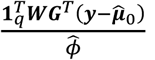 are asymptotically normal, and their covariance matrix is

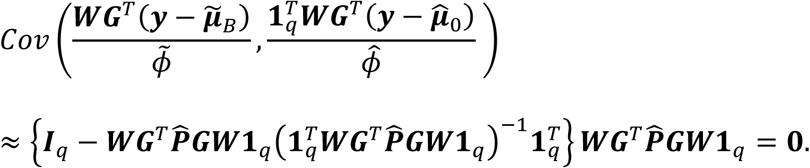

Therefore, *T*_*θ*_ and *T*_*B*_ are approximately asymptotically independent. Let *p*_*θ*_ and *p*_*B*_ be the p value of the two tests respectively, then SMMAT-E p value *p*_*E*_ is computed using Fisher’s method with a chi-square distribution with 4 df as 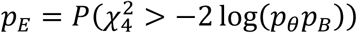.

### Meta-analysis

SMMAT-B, SMMAT-S, SMMAT-O and SMMAT-E can all be conducted in the meta-analysis context. Assuming the single-variant scores 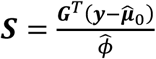 and their covariance matrix 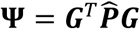 are computed for each variant set in each study, we can reconstruct 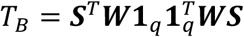 with 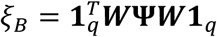; *T*_*S*_ = ***S***^T^***WWS*** with **Ξ**_*S*_ *=* ***WΨW***; *T*_*ρ*_ *= ρT*_*B*_+ (1 - *ρ*)*T*_*S*_and 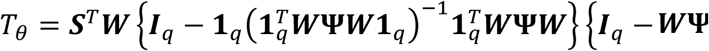 with 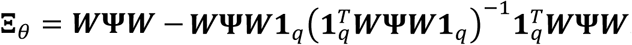

For each variant set, let *m* = 1, 2, …, *M* be the index of studies, ***S***_*m*_ and **Ψ**_*m*_ be the single-variant scores and covariance matrix from study *m*, in testing the “weak” null hypothesis^26^ of summary genetic effects *H*_0_: ***β*** = **0**,^24, 25^ we can compute meta summary statistics 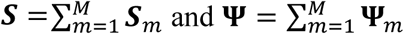 and use them in SMMAT-B, SMMAT-S, SMMAT-O and SMMAT-E. When combining studies with very different sample characteristics, testing the “strong” null hypothesis^26^ that genetic effects in all studies are 0 is sometimes desired. In the general case, we may choose to group studies that are similar and test if the summary genetic effects in all groups are 0, for example, in the meta-analysis of multi-ethnic samples. Let *c* = 1, 2, …, *C* be a partition of *M* studies (*C* ≤ *M*), where *C* is the number of ethnicities, 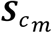 and 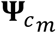 be the single-variant scores and covariance matrix from study *m* in partition *c* (*m* = 1, 2, …, *M*_*c*_ in partition *c*, and 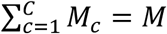), such that genetic effects for the same variant are summarized within each partition *c* but heterogeneous across partitions,^24^ we can also compute summary statistics 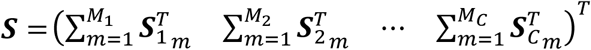 and 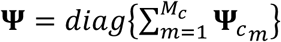. Note that ***S*** is now a vector of length *Cq*, and **Ψ** is a block-diagonal matrix with *C* blocks of *q* × *q* matrices, one for each partition of studies (with total dimension *Cq* × *Cq*), we should replace ***W***, **1**_*q*_ and ***I***_*q*_ by ***I***_*C*_ ⊗ ***W*** (where ⊗ denotes the Kronecker product), **1**_*Cq*_ and ***I***_*Cq*_, respectively in the above expressions for *T*_*B*_, *T*_*ρ*_, *T*_*S*_ and *T*_*θ*_ for meta-analysis.

### Simulation studies

#### Type I error in single-cohort studies

We performed coalescent simulations to generate sequence data with 100 genetic variants in each set, and 10,000 independent sets for 8,000 individuals from a 20 × 20 grid of spatially continuous populations with migration rate between adjacent cells *M* = 10 (Figure 1A). Within each cell, we paired 20 individuals into 10 families and simulated 2 children for each family using gene dropping,^27^ and in total we had 4,000 families and 16,000 individuals. For continuous traits, in each simulation replicate, we simulated the phenotype *y*_*ij*_ for individual *j* in family *i* under the null hypothesis of no genetic association from

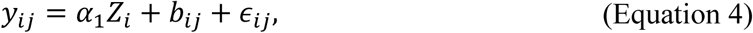

**Figure 1.**
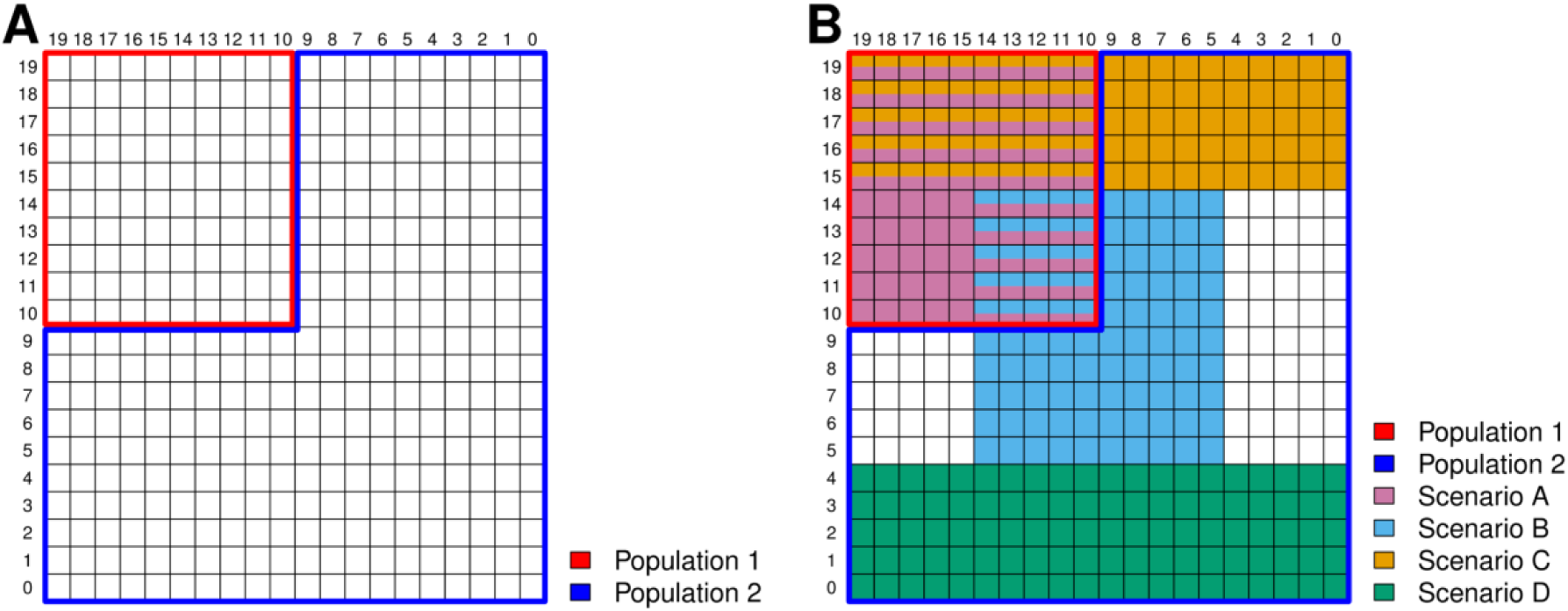
Map of spatially continuous populations from which genotypes were simulated based on the coalescent model. (A) Map for a single-cohort simulation study: the top left 10 × 10 grid formed Population 1, and the rest formed Population 2. (B) Map for a meta-analysis simulation study: Scenario A studies were unrelated individuals sampled from Population 1 only; Scenario B studies were related individuals sampled from specific regions in Population 1 and Population 2; Scenario C studies were unrelated individuals sampled from specific regions in Population 1 and Population 2; and Scenario D studies were related individuals sampled from specific regions in Population 2 only.

where the “population effect” *α*_1_ = 1, the population indicator *Z*_*i*_ = 1 if family *i* was from a 10 × 10 grid in the top left of the map (Population 1), and *Z*_*i*_= 0 otherwise (Population 2). The familial random effects were simulated as

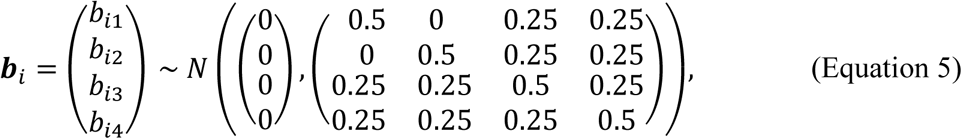

and the random error *∈*_*ij*_ ∼ *N*(0, 1) for each individual *j* in family *i*. Then we randomly sampled 3,500 individuals from the 10 × 10 grid in the top left, and 6,500 individuals from the rest of the map. The family identifier was removed for all individuals in the analysis, so that there were both population structure and cryptic relatedness in the sample. We compared SMMAT-B, SMMAT-S, SMMAT-O and SMMAT-E in analyzing 10,000 independent variant sets based on a linear mixed model using our GMMAT package, including random effects with their covariance matrix proportional to the GRM, and adjusted for the first 10 principal components (PCs) of ancestry. We repeated this 4,000 times to get p values combined from 40 million independent genetic variant sets for each test.

For binary traits, in each simulation replicate, we simulated the phenotype *y*_*ij*_ for individual *j* in family *i* under the null hypothesis of no genetic association from

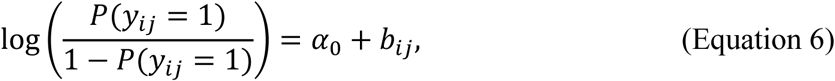

where *α*_0_ was chosen such that the disease prevalence was 0.01 in all populations, and the familial random effects *b*_*ij*_ were simulated in the same way as for continuous traits. Then we randomly sampled 2,500 cases and 1,000 controls from the 10 × 10 grid in the top left (Population 1), and 2,500 cases and 4,000 controls from the rest of the map (Population 2) to form a hypothetical study with balanced cases and controls in combined populations. Therefore, there was confounding by population structure resulting from unequal sampling, even though the disease prevalence was the same. We removed the family identifier, compared SMMAT-B, SMMAT-S, SMMAT-O and SMMAT-E in analyzing 10,000 independent variant sets based on a logistic mixed model using our GMMAT package, similarly as described above, and repeated this 4,000 times to get p values combined from 40 million independent genetic variant sets for each test.

#### Type I error in meta-analysis

We also conducted simulation studies in the meta-analysis context to evaluate the type I error rates. We considered 4 scenarios: unrelated individuals, without confounding by population structure (Scenario A studies); related individuals, with confounding by population structure (Scenario B studies); unrelated individuals, with confounding by population structure (Scenario C studies); and related individuals, without confounding by population structure (Scenario D studies).

For Scenario A studies, we simulated 16 unrelated individuals in each cell from the 10 × 10 grid in the top left of the map (Figure 1B). For continuous traits, we simulated the phenotype *y*_*ij*_ from Equation 4, with *α*_1_ = 0 and *b*_*ij*_ = 0, and randomly sampled 1,000 individuals. For binary traits, we simulated *y*_*ij*_ from Equation 6, with *b*_*ij*_ = 0, and randomly sampled 500 cases and 500 controls.

For Scenario B studies, we simulated 8 unrelated individuals, paired them into 4 families and simulated 2 children for each family in each cell from the 10 × 10 grid in the center of the map (Figure 1B). For continuous traits, we simulated the phenotype *y*_*ij*_ from Equation 4, with *α*_1_ = 1, the population indicator *Z*_*i*_ = 1 if family *i* was from Population 1, and *Z*_*i*_ = 0 if from Population 2, and familial random effects *b*_*ij*_ were simulated using Equation 5, and we randomly sampled 350 individuals from Population 1 and 650 individuals from Population 2. For binary traits, we simulated *y*_*ij*_ from Equation 6, with *b*_*ij*_ from Equation 5, and randomly sampled 250 cases and 100 controls from Population 1, and 250 cases and 400 controls from Population 2.

For Scenario C studies, we simulated 16 unrelated individuals in each cell from the 20 × 5 grid in the top of the map (Figure 1B). For continuous traits, we simulated the phenotype *y*_*ij*_ from Equation 4, with *α*_1_ = 1, the population indicator *Z*_*i*_ = 1 if family *i* was from Population 1, and *Z*_*i*_ = 0 if from Population 2, and *b*_*ij*_ = 0, and we randomly sampled 350 individuals from Population 1 and 650 individuals from Population 2. For binary traits, we simulated *y*_*ij*_ from Equation 6, with *b*_*ij*_ = 0, and randomly sampled 250 cases and 100 controls from Population 1, and 250 cases and 400 controls from Population 2.

For Scenario D studies, we simulated 8 unrelated individuals, paired them into 4 families and simulated 2 children for each family in each cell from the 20 × 5 grid in the bottom of the map (Figure 1B). For continuous traits, we simulated the phenotype *y*_*ij*_ from Equation 4, with *α*_1_ = 0, familial random effects *b*_*ij*_ simulated using Equation 5, and we randomly sampled 1,000 individuals. For binary traits, we simulated *y*_*ij*_ from Equation 6, with *b*_*ij*_ from Equation 5, and randomly sampled 500 cases and 500 controls.

In each simulation replicate, we simulated 3 studies from each scenario, totaling 12 studies with a combined sample size of 12,000 (6,000 cases and 6,000 controls for binary traits). We compared SMMAT-B, SMMAT-S, SMMAT-O and SMMAT-E using two meta-analysis strategies: all studies in the same group, and Scenario A, B, C, D studies in 4 separate groups. In the latter case, 3 studies from the same scenario were grouped in the same partition with shared genetic effects, while studies from different scenarios were allowed to have heterogeneous genetic effects. We repeated 4,000 simulation replicates to get p values from 40 million independent genetic variant sets.

#### Power

We used the same genotype data as in the single-cohort type I error simulations and evaluated the empirical power of SMMAT-B, SMMAT-S, SMMAT-O and SMMAT-E (with weights equal to a beta distribution density function with parameters 1 and 25 on the MAF of each variant^13^) in 9 scenarios, with the proportion of causal variants in a test unit ranging from 10%, 20% to 50%, and the proportion of variants with negative effects out of causal variants ranging from 100%, 80% to 50%. For continuous traits, we simulated the phenotype *y*_*ij*_ for individual *j* in family *i* from

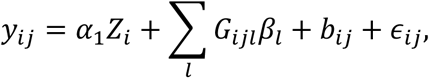

where *α*_1_ = 1, the population indicator *Z*_*i*_ = 1 if family *i* was from Population 1, and *Z*_*i*_ = 0 if from Population 2, *g*_*ijl*_ was the centered genotype for causal variant *l* of individual *j* in family *i*, the causal effect size was |*β*_*l*_| = *c*|log_10_ *MAF*_*l*_| for variant *l* with *MAF*_*l*_, where the constant *c* was set to 0.2, 0.1 and 0.05 when the proportion of causal variants was 10%, 20% and 50%, the familial random effects *b*_*ij*_ were simulated using Equation 5, and the random error *∈*_*ij*_ ∼ *N*(0, 1). We randomly sampled 35% individuals from Population 1, and 65% individuals from Population 2.

For binary traits, we simulated the phenotype *y*_*ij*_ for individual *j* in family *i* from

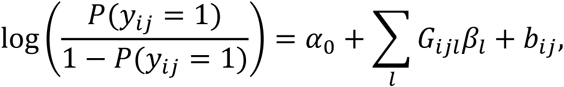

where *α*_0_ was chosen such that the disease prevalence was 0.01 in all populations, *G*_*ijl*_ was the centered genotype for causal variant *l* of individual *j* in family *i*, the causal effect size was |*β*_*l*_| = *c*|log_10_ *MAF*_*l*_| for variant *l* with *MAF*_*l*_, where the constant *c* was set to 0.3, 0.2 and 0.1 when the proportion of causal variants was 10%, 20% and 50%, the familial random effects *b*_*ij*_ were simulated using Equation 5. We randomly sampled 35% individuals (with 25% cases and 10% controls out of the total sample size) from Population 1, and 65% individuals (with 25% cases and 40% controls out of the total sample size) from Population 2 to form a hypothetical study with balanced cases and controls in combined populations.

For both continuous and binary traits, we varied the total sample size from 2,000, 5,000 to 10,000, repeated 1,000 simulation replicates for each scenario under the alternative hypothesis, and compared the empirical power at the significance level of 2.5 × 10^−6^.

### TOPMed example involving fibrinogen levels

Samples with both plasma fibrinogen measures and whole genome sequence data (Freeze 5b) from the following 11 TOPMed studies were included in the analysis: the Old Order Amish Study (Amish), Cleveland Family Study (CFS), Genetic Epidemiology of COPD Study (COPDGene), Framingham Heart Study (FHS), Jackson Heart Study (JHS), San Antonio Family Study (SAFS), the Atherosclerosis Risk in Communities (ARIC) Study, Genetic Studies of Atherosclerosis Risk (GeneSTAR), Genetic Epidemiology Network of Arteriopathy (GENOA), the Multi-Ethnic Study of Atherosclerosis (MESA), and Women’s Health Initiative (WHI). The TOPMed studies were approved by institutional review boards at participating institutions, and informed consent was obtained from all study participants. Amish, CFS, FHS, JHS, and SAFS are family-based studies with differing degrees of relatedness. The total sample size was 23,763. Within each study and each ethnicity, measured fibrinogen levels were adjusted for age, sex, study-specific covariates, and the residuals were rank normalized and rescaled by multiplying by the original standard deviation, so that the transformed phenotype data have the same variances as on the original scale. The transformed phenotype data were pooled together in the analysis, using a heteroscedastic linear mixed model^28^ allowing for different residual variances in each study/ethnicity, adjusting for study, ethnicity, sequence center, top 10 ancestry PCs^29^ as fixed-effects covariates, and including a GRM calculated by Mixed Model Analysis for Pedigrees and Populations (MMAP) to model the random effects for relatedness. Rare and low frequency genetic variants on chromosome 4 with MAF less than 5% were tested for association with fibrinogen levels in a sliding window analysis^30^ of 4 kb non-overlapping windows, using SMMAT-B, SMMAT-S, SMMAT-O and SMMAT-E with weights equal to a beta distribution density function with parameters 1 and 25 on the MAF of each variant^13^. The analysis was performed using the GMMAT App (version 0.9.2) with 32 parallel threads on a single computing node with 240 GB total memory in the Analysis Commons.^31^ To benchmark the computational speed in running SMMAT-B, SMMAT-S, SMMAT-O and SMMAT-E, we also ran re-analyses to perform each test separately, using summary statistics from the sliding window analysis and a single thread on a computing node with 15 GB total memory in the Analysis Commons.

## RESULTS

### Simulation studies

Table 1 shows the empirical type I error rates of SMMAT-B, SMMAT-S, SMMAT-O and SMMAT-E at significance levels of 0.05, 0.0001, and 2.5 × 10^−6^, in the variant set analyses of continuous and binary traits in single-cohort simulation studies. All 4 tests have well-controlled type I error rates at these significance levels, suggesting that GLMMs can be effective in adjusting for population structure and cryptic relatedness in complex study samples. This is also consistent with the quantile-quantile (QQ) plots in Figure 2, which show neither inflation nor deflation in the tail.

**Table 1.** Empirical type I error rates of SMMAT-B, SMMAT-S, SMMAT-O and SMMAT-E in single-cohort simulation studies at significance levels of 0.05, 0.0001, and 2.5 × 10^−6^. The total sample size was 10,000, and results from 4,000 simulation replicates were combined to get 40 million genetic variant sets.

| Level | Continuous Traits |  |  | Binary Traits |  |  |
| --- | --- | --- | --- | --- | --- | --- |
| | 0.05 | 0.0001 | $2.5 \times 10^{-6}$ | 0.05 | 0.0001 | $2.5 \times 10^{-6}$ |
| SMMAT-B | 0.047 | $8.7 \times 10^{-5}$ | $2.0 \times 10^{-6}$ | 0.049 | $9.6 \times 10^{-5}$ | $2.0 \times 10^{-6}$ |
| SMMAT-S | 0.048 | $8.7 \times 10^{-5}$ | $2.0 \times 10^{-6}$ | 0.049 | $9.5 \times 10^{-5}$ | $2.3 \times 10^{-6}$ |
| SMMAT-O | 0.050 | $1.1 \times 10^{-4}$ | $3.0 \times 10^{-6}$ | 0.052 | $1.2 \times 10^{-4}$ | $3.0 \times 10^{-6}$ |
| SMMAT-E | 0.050 | $1.0 \times 10^{-4}$ | $3.0 \times 10^{-6}$ | 0.050 | $9.9 \times 10^{-5}$ | $2.0 \times 10^{-6}$ |

**Figure 2.**
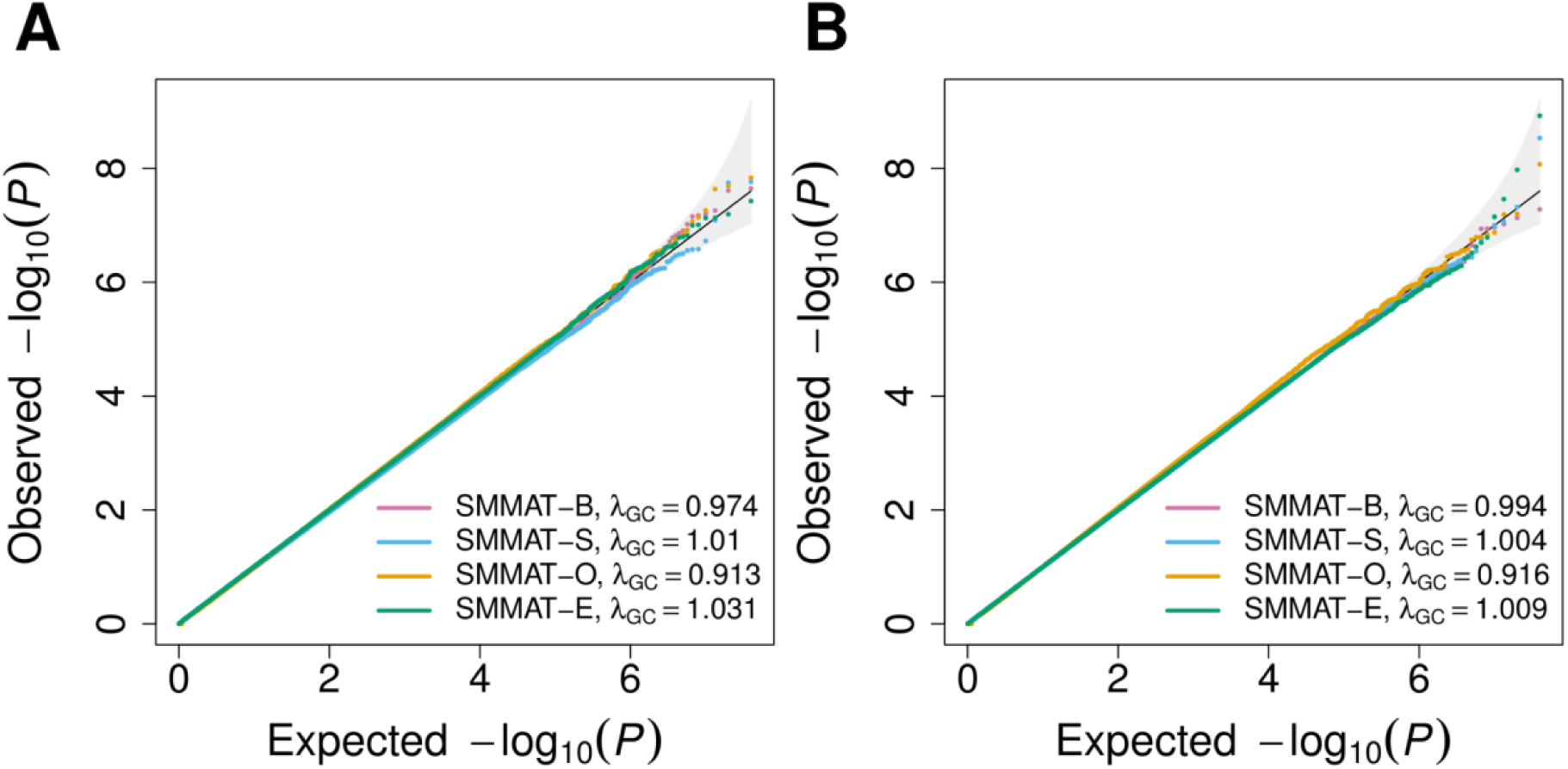
Quantile-quantile plots of SMMAT-B, SMMAT-S, SMMAT-O and SMMAT-E in the analysis of 10,000 samples in single-cohort studies with both population structure and cryptic relatedness, under the null hypothesis of no genetic association. (A) Continuous traits in linear mixed models. (B) Binary traits in logistic mixed models.

Table 2 and Figure 3 show simulation results of SMMAT-B, SMMAT-S, SMMAT-O and SMMAT-E assuming all studies in the same group (hom) or in 4 separate groups (het) in meta-analyses for combining 4 types of studies: with and without confounding by population structure, with and without cryptic relatedness. We note that SMMAT-B statistic *T*_*B*_ has the same form in these two meta-analysis strategies,^24^ therefore, we included 7 tests in the simulation studies. In het SMMAT-S, SMMAT-O and SMMAT-E, studies from the same scenario were grouped together to assume shared genetic effects. Under the null hypothesis of no genetic associations, hom SMMAT-O shows very mild inflation in our simulation settings, but all other 6 tests in the SMMAT framework control type I error rates well at significance levels of 0.05, 0.0001, and 2.5 × 10^−6^ and have well-calibrated tail probabilities, for both continuous and binary traits.

**Table 2.** Empirical type I error rates of SMMAT-B, SMMAT-S, SMMAT-O and SMMAT-E assuming all studies in the same group (hom) and Scenario A, B, C, D studies in 4 separate groups (het), in meta-analysis simulation studies at significance levels of 0.05, 0.0001, and 2.5 × 10^−6^. The total sample size was 12,000 from 12 studies, and results from 4,000 simulation replicates were combined to get 40 million genetic variant sets.

| Level | Continuous Traits |  |  | Binary Traits |  |  |
| --- | --- | --- | --- | --- | --- | --- |
| | 0.05 | 0.0001 | $2.5 \times 10^{-6}$ | 0.05 | 0.0001 | $2.5 \times 10^{-6}$ |
| SMMAT-B | 0.051 | $1.0 \times 10^{-4}$ | $2.6 \times 10^{-6}$ | 0.051 | $1.1 \times 10^{-4}$ | $2.5 \times 10^{-6}$ |
| Hom SMMAT-S | 0.051 | $1.0 \times 10^{-4}$ | $2.6 \times 10^{-6}$ | 0.051 | $1.1 \times 10^{-4}$ | $2.1 \times 10^{-6}$ |
| Het SMMAT-S | 0.051 | $1.0 \times 10^{-4}$ | $2.8 \times 10^{-6}$ | 0.052 | $1.0 \times 10^{-4}$ | $2.4 \times 10^{-6}$ |
| Hom SMMAT-O | 0.053 | $1.3 \times 10^{-4}$ | $4.0 \times 10^{-6}$ | 0.053 | $1.4 \times 10^{-4}$ | $3.4 \times 10^{-6}$ |
| Het SMMAT-O | 0.052 | $1.1 \times 10^{-4}$ | $2.6 \times 10^{-6}$ | 0.052 | $1.1 \times 10^{-4}$ | $2.2 \times 10^{-6}$ |
| Hom SMMAT-E | 0.051 | $1.0 \times 10^{-4}$ | $2.5 \times 10^{-6}$ | 0.051 | $1.1 \times 10^{-4}$ | $2.6 \times 10^{-6}$ |
| Het SMMAT-E | 0.051 | $1.0 \times 10^{-4}$ | $2.8 \times 10^{-6}$ | 0.052 | $1.1 \times 10^{-4}$ | $3.0 \times 10^{-6}$ |

**Figure 3.**
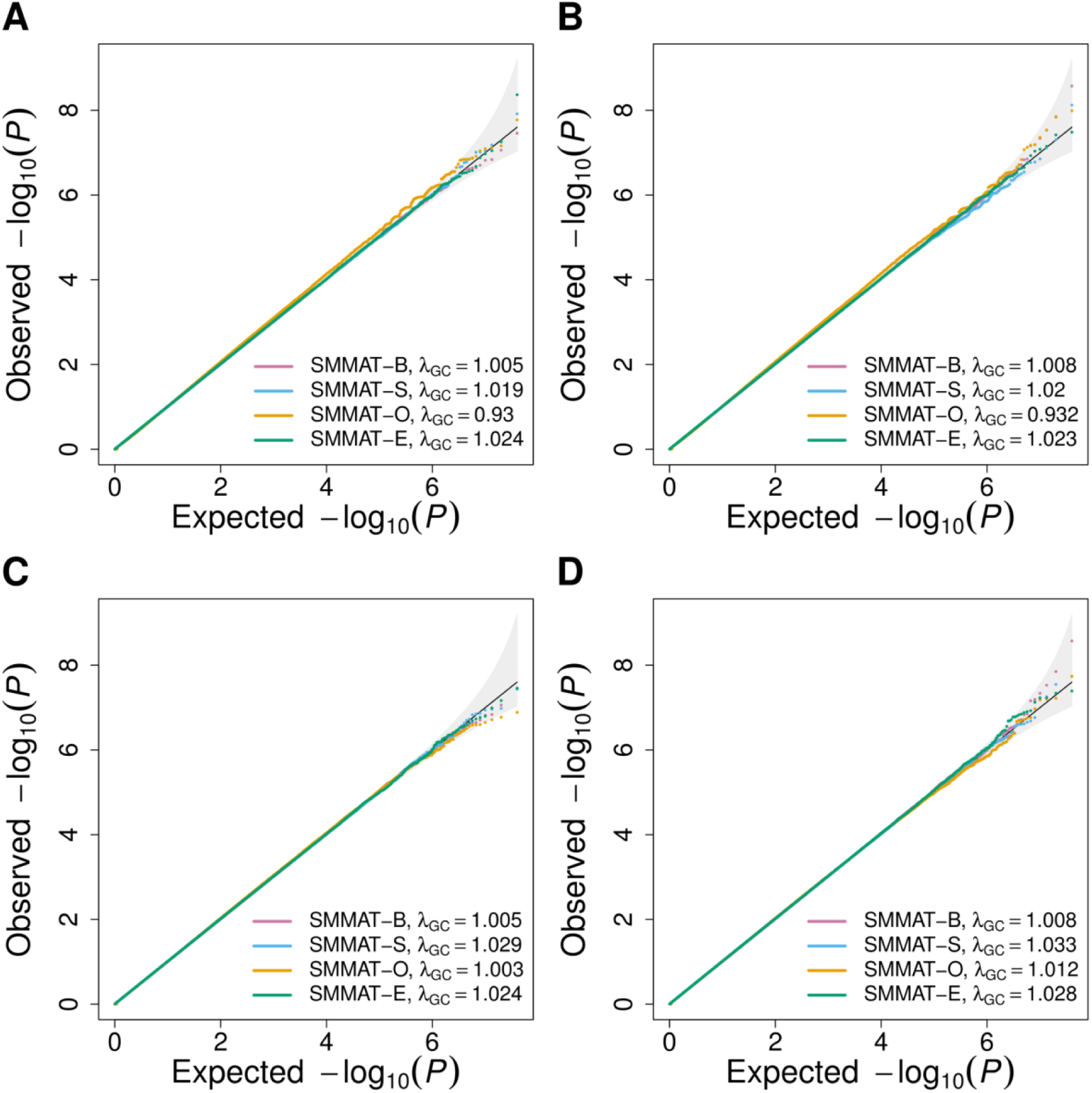
Quantile-quantile plots of SMMAT-B, SMMAT-S, SMMAT-O and SMMAT-E in the meta-analysis of 12 studies with a total sample size of 12,000, under the null hypothesis of no genetic association. (A) Continuous traits in linear mixed models, all studies in the same group. (B) Binary traits in logistic mixed models, all studies in the same group. (C) Continuous traits in linear mixed models, Scenario A, B, C, D studies in 4 separate groups. (D) Binary traits in logistic mixed models, Scenario A, B, C, D studies in 4 separate groups.

Figures 4 and 5 present the empirical power for causal variant sets at the significance level of 2.5 × 10^−6^ for continuous and binary traits, respectively. The power increases with the sample size. As the proportion of causal variants with effects in the same direction drops from 100%, 80% to 50% in each row, the power drops for all tests, but most substantially for the burden test SMMAT-B. When the sample size is large (i.e., 10,000 samples), SMMAT-E has the highest power, for both continuous and binary traits in all 9 simulation scenarios.

**Figure 4.**
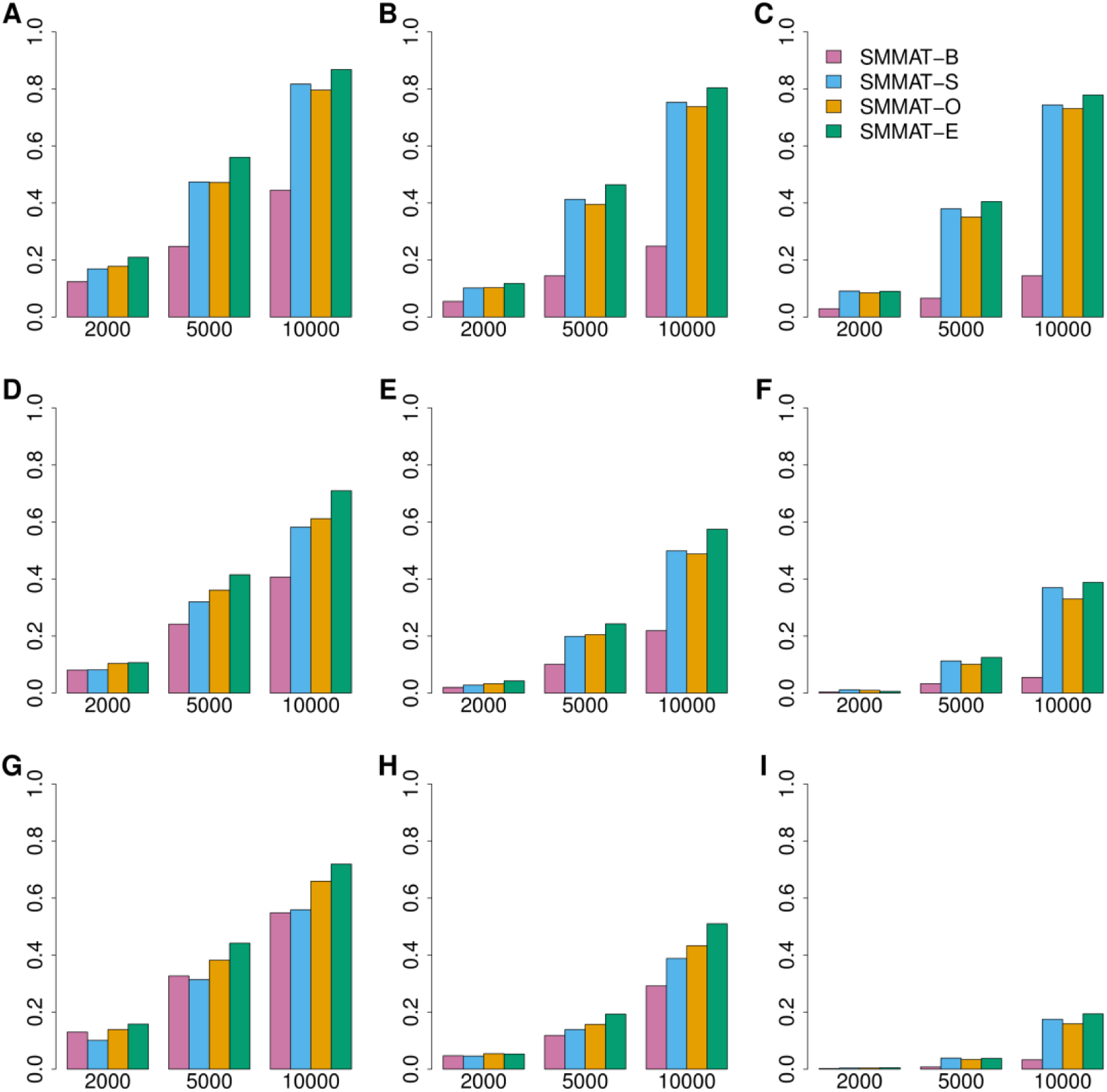
Empirical power of linear mixed model based SMMAT-B, SMMAT-S, SMMAT-O and SMMAT-E in continuous trait analysis of 2,000, 5,000 and 10,000 samples. (A) 10% causal variants with 100% negative effects. (B) 10% causal variants with 80% negative effects. (C) 10% causal variants with 50% negative effects. (D) 20% causal variants with 100% negative effects. (E) 20% causal variants with 80% negative effects. (F) 20% causal variants with 50% negative effects. (G) 50% causal variants with 100% negative effects. (H) 50% causal variants with 80% negative effects. (I) 50% causal variants with 50% negative effects. Effect sizes were simulated using the same parameter in each row, but different across rows.

**Figure 5.**
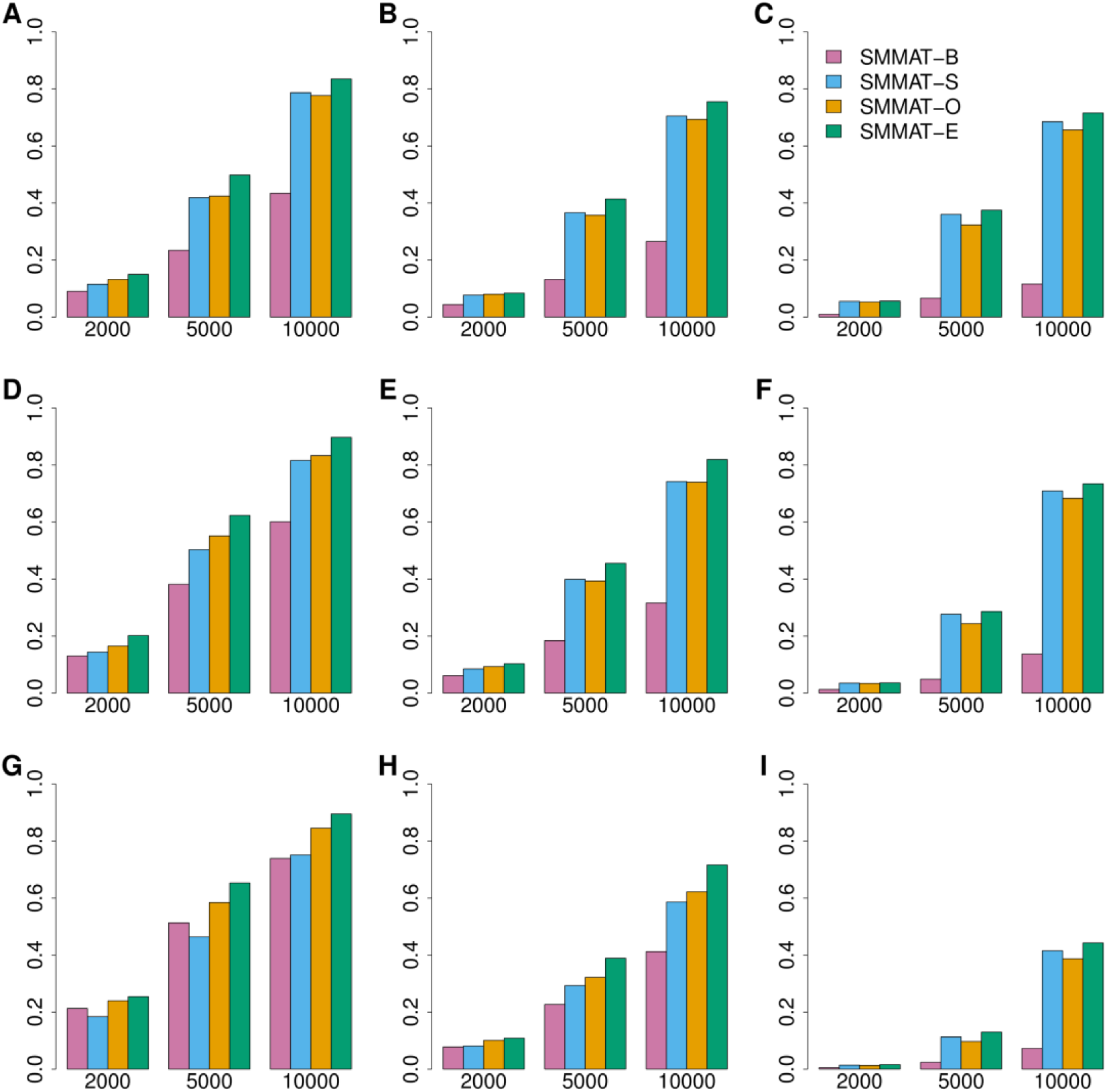
Empirical power of logistic mixed model based SMMAT-B, SMMAT-S, SMMAT-O and SMMAT-E in binary trait analysis of 2,000, 5,000 and 10,000 samples. (A) 10% causal variants with 100% negative effects. (B) 10% causal variants with 80% negative effects. (C) 10% causal variants with 50% negative effects. (D) 20% causal variants with 100% negative effects. (E) 20% causal variants with 80% negative effects. (F) 20% causal variants with 50% negative effects. (G) 50% causal variants with 100% negative effects. (H) 50% causal variants with 80% negative effects. (I) 50% causal variants with 50% negative effects. Effect sizes were simulated using the same parameter in each row, but different across rows.

### TOPMed example involving fibrinogen levels

We compared the results from SMMAT-B, SMMAT-S, SMMAT-O and SMMAT-E in an analysis of fibrinogen levels, using chromosome 4 (including the genomic region that encodes the fibrinogen protein, *FGB*) whole genome sequence data from 11 TOPMed studies. Previous studies have reported two rare variants within *FGB* on chromosome 4, rs6054 (hg38 position 154,568,456) and rs201909029 (hg 38 position 154,567,636) associated with lower fibrinogen levels, with similar effect sizes in all ancestry groups.^32^ In the sliding window analysis, we grouped low frequency and rare genetic variants with MAF less than 5% into 46,859 non-overlapping 4 kb windows containing at least one variant. The number of variants in each window passing the MAF filter ranged from 1 to 1,290, with a median of 351 (25% quartile 326 and 75% quartile 380). The QQ plot (Figure 6A) shows that all 4 tests have well-calibrated tail probabilities. Table 3 summarizes heteroscedastic linear mixed model-based SMMAT-B, SMMAT-S, SMMAT-O and SMMAT-E p values in *FGB* and flanking regions. SMMAT-S, SMMAT-O and SMMAT-E give the most significant results in the 4 kb window 154,554 – 154,558 kb, with p values 1.6 × 10^−17^, 8.9 × 10^−17^, and 6.2 × 10^−19^, respectively, while SMMAT-B p value is much larger (6.9 × 10^−5^). In the 4 kb window that covers both known association rare variants rs6054 and rs201909029 (window 154,566 – 154,570 kb), SMMAT-E gives the smallest p value 3.1 × 10^−17^), followed by SMMAT-S (p value 9.7 × 10^−17^), SMMAT-O (p value 3.3 × 10^−16^) and SMMAT-B (p value 1.6 × 10^−8^).

**Figure 6.**
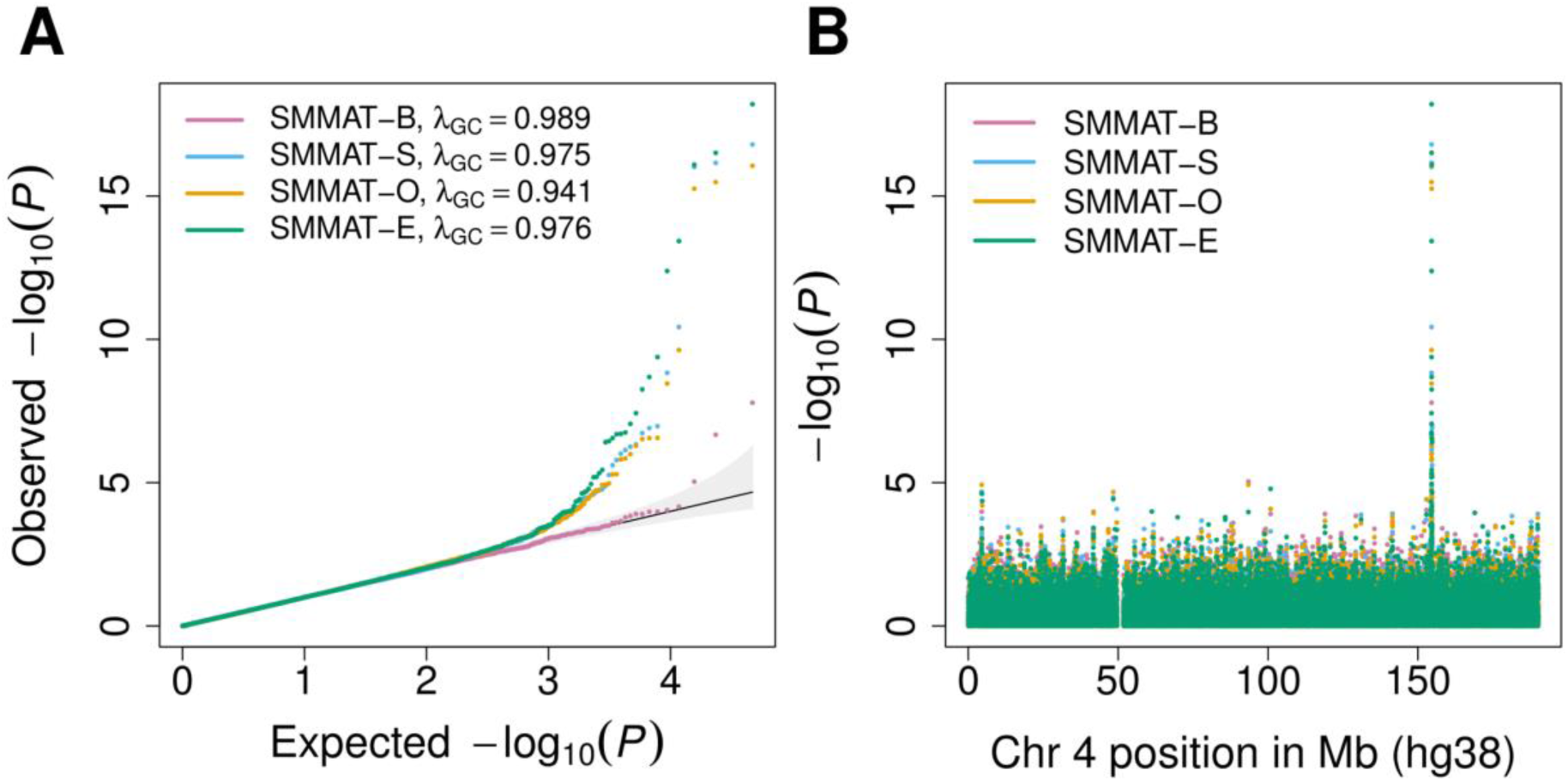
TOPMed fibrinogen level SMMAT analysis results using a heteroscedastic linear mixed model on rare variants with MAF < 5% in non-overlapping 4 kb sliding windows on chromosome 4 (n = 23,763). (A) Quantile-quantile plot. (B) P values on the log scale versus physical positions of the windows on chromosome 4 (build hg38).

**Table 3.**
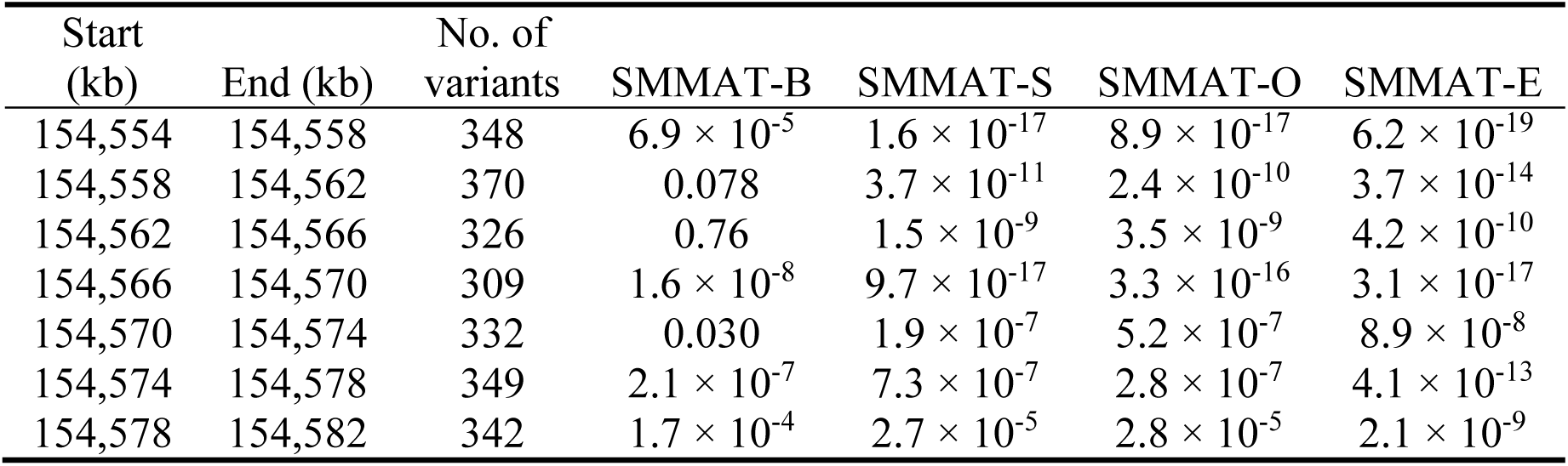
TOPMed fibrinogen level SMMAT p values in known association gene *FGB* and flanking regions on chromosome 4, using a heteroscedastic linear mixed model on rare variants with MAF < 5% (n = 23,763). Physical positions of each window are on build hg38.

### Computation time

Table 4 shows the CPU time for running the sliding window analysis for 23,763 individuals with TOPMed whole genome sequence data and fibrinogen levels, using summary statistics from 46,859 non-overlapping 4 kb windows on chromosome 4. The GMMAT App (version 0.9.2) in the Analysis Commons cloud computing platform has implemented SMMAT-B, SMMAT-S, SMMAT-O and SMMAT-E, with the option of running one or more tests in an analysis. SMMAT-B results are automatically included when running SMMAT-O or SMMAT-E, and SMMAT-S p values will also be output when running SMMAT-O. Of the four tests in Table 4, SMMAT-B takes shortest time as the p value calculation does not involve any eigen-decomposition of covariance matrices. SMMAT-S takes only about 10 minutes longer than SMMAT-B for the eigen-decomposition of 46,859 covariance matrices. SMMAT-E takes about 12 minutes longer than SMMAT-S and gives both SMMAT-B and SMMAT-E p values. SMMAT-O takes 175 minutes longer than SMMAT-S, as more eigen-decompositions are performed in SMMAT-O when it searches for the optimal combination of SMMAT-B and SMMAT-S on a grid of *ρ* values.

**Table 4.** CPU time in the TOPMed fibrinogen level SMMAT using summary statistics from a sliding window analysis using non-overlapping 4 kb windows on chromosome 4 (n = 23,763). Tests were performed using the GMMAT App (version 0.9.2) with one single thread on a computing node with 15 GB total memory in the Analysis Commons.

| Test | Time (min) |
| --- | --- |
| SMMAT-B | 81 |
| SMMAT-S | 91 |
| SMMAT-O | 266 |
| SMMAT-E | 103 |

## DISCUSSION

We have developed and implemented SMMAT, a family of computationally-efficient variant set mixed model association tests for continuous and binary traits in large-scale whole genome sequencing studies. This framework includes extensions of three widely used variant set tests for unrelated individuals to complex study samples with population structure and cryptic relatedness: the burden test (SMMAT-B), SKAT (SMMAT-S) and SKAT-O (SMMAT-O), as well as a new efficient hybrid test that combines the mixed model burden and SKAT tests (SMMAT-E). Specifically, SMMAT-E is constructed by combining the burden test and an adjusted mixed model SKAT statistic that is approximately asymptotically independent from the mixed model burden test statistic, in a similar spirit to MiST in non-mixed model setting,^15^ but that differs from MiST in that it does not require fitting separate mixed effect burden models for each variant set with the set genetic burden as a fixed-effects covariate. Instead, we use matrix projections to approximate the adjusted SKAT statistic from a global null model without any fixed effects for the variant set-specific genetic burden. Of note, this global null model only needs to be fit once in a whole genome analysis, which greatly reduces the computational cost. We show in simulation studies and the TOPMed fibrinogen example that SMMAT-E is more powerful than the other three tests in large samples, at the computational cost almost on the same scale of SMMAT-B and SMMAT-S. Therefore, SMMAT-E is recommended in the analysis of large-scale whole genome sequencing studies.

In the SMMAT framework, different weighting strategies can be used. One can use a function of the MAF,^11, 13^ or external measures based on functional annotation such as CADD,^33^ Eigen,^34^ FATHMM-XF,^35^ or tissue-specific annotations, such as GENOSKYLINE,^36^ as the weight for each variant in a set. In the analysis of fibrinogen levels in TOPMed, we used MAF-based weights. Recently, unified variant set tests allowing for multiple functional annotations have been developed,^37^ and the SMMAT framework can possibly be extended to accommodate multiple weights. Nevertheless, the optimal weighting strategy in rare variant analysis remains an open question and an active field of research.

As SMMAT-E combines the burden test p value *p*_*B*_ with an asymptotically independent adjusted SKAT p value *p*_*θ*_ using Fisher’s method in our SMMAT implementation in the GMMAT App, we note that other forms of combinations may also be applied.^38^ For example, previous studies have shown that Tippett’s procedure based on the minimum of *p*_*θ*_ and *p*_*B*_ might be more powerful than Fisher’s method in MiST when only one of the p values is small.^15^ Alternatively, instead of combining the p values, weighted linear combinations of chi-square statistics have been proposed,^39-41^ and they can also be applied to combine the burden test statistic *T*_*B*_ and the asymptotically independent SKAT statistic *T*_*θ*_ in the SMMAT framework.

SMMAT also has some limitations. SMMAT p values are computed based on asymptotic distributions, which may be not be accurate in small samples, especially for binary traits and heavily skewed continuous traits. For continuous traits, small-sample inference procedures have been proposed for SKAT,^42, 43^ and the same methodology can be applied to SMMAT. For ultra-rare genetic variants with very low minor allele counts, the single-variant scores used to construct SMMAT-B, SMMAT-S, SMMAT-O and SMMAT-E may not be close to a normal distribution, even if the total sample size is large. If there are only ultra-rare variants (e.g. singletons, doubletons) in a test region and the number of variants is small, SMMAT-B might be the best analysis strategy as its asymptotic property depends on the cumulative minor allele counts. Moreover, the asymptotic issue of single-variant scores also exists for binary traits with highly unbalanced case-control ratios, and a saddlepoint approximation approach has been proposed to match the cumulant generating function of the single-variant scores,^44^ and it has recently been extended to GLMMs.^45^

Fitting GLMMs with a GRM has *O*(*n*^3^) complexity in general, where *n* is the sample size. We have overcome this computational challenge by fitting only one GLMM in a whole genome analysis, and using matrix multiplications with *O*(*n*^2^) complexity for each variant set in SMMAT. In large-scale whole genome sequencing studies, solutions to other computational challenges are being proposed. For example, when the number of variants *q* in SKAT is very large, eigendecomposition of the covariance matrix, which has *O*(min(*n*, *q*)^3^) complexity, could be computationally expensive. Recently, the fastSKAT approach has been proposed to efficiently approximate the null distribution of SKAT when *q* is very large,^46^ and the same strategy can be applied to speed up SMMAT p value calculation for very large *q*. On the other hand, as the sample size in ongoing large-scale sequencing projects such as TOPMed eventually expands to hundreds of thousands, using a full *n* × *n* GRM would not be computationally practical in pooled analyses, as it may take several weeks to fit even only one GLMM with *O*(*n*^3^) complexity, and *O*(*n*^2^) memory footprint. Meta-analyses may be a more appealing analysis strategy in that situation by combining summary statistics from study-specific or ancestry-specific analyses. Essentially equivalently, in pooled analyses, using a sparse and/or block-diagonal GRM with each block corresponding to an individual study in meta-analyses, will help reduce the computational cost in fitting GLMMs, providing one uses specialized routines for manipulation of sparse matrices.^47^ Although whole genome sequencing studies have not yet been conducted in large biobanks with sample sizes on the scale of millions of individuals, it is expected that calculating the GRM itself would become a major computational bottleneck. Recently, GRM-free mixed effects models such as BOLT-LMM^6, 48^ and SAIGE^45^ have been developed for single variant tests, and we note that extension of these methods to the SMMAT framework will further reduce the computational cost in biobank-scale whole genome sequencing studies in the future.

In summary, SMMAT provides a flexible and practical statistical framework for large-scale whole genome sequencing studies with complex study samples, with balanced power and computational performance. With continuing advances in technology, lowering cost and development of new analytical methods, large-scale whole genome sequencing studies will facilitate human genetic research and enhance our understandings on complex diseases and traits.

## Appendix: Approximations in SMMAT-E

Here we derive the approximations used in SMMAT-E to construct the SKAT-type statistic adjusting for the genetic burden

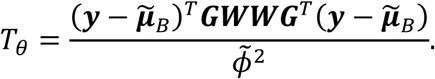

Let 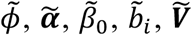 and 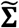 be estimates for *Φ*, ***α***, *β*, *b*, ***V*** and **∑** respectively from the burden GLMM (Equation 3), we define 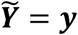 as the phenotype vector for continuous traits, and the “working vector” with components 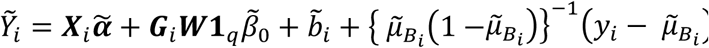 at convergence of the logistic burden mixed model for binary traits (Equation 3), where 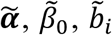 are fixed-effects and random-effects estimates from the burden GLMM. We have

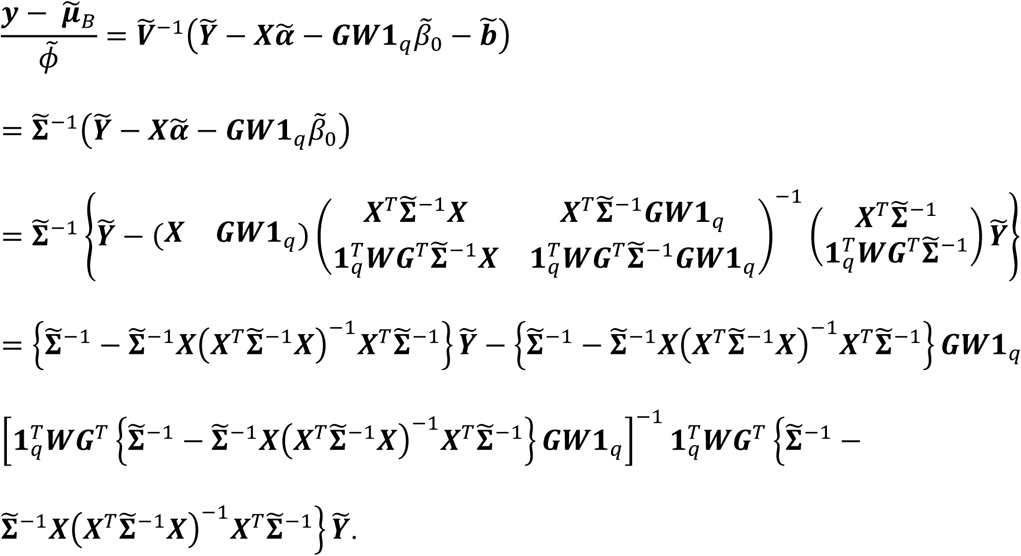

Note that 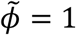 for binary traits. Moreover, since the true value of *β*_0_ is small, assuming including the genetic burden ***G***_*i*_***W*1**_*q*_ in the second term in Equation 3 does not dramatically change the variance component estimates for *τ*_*k*_ and *Φ* (and for binary traits, also the “working vector” 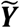 at convergence of the model from Equation 2), we have the approximation 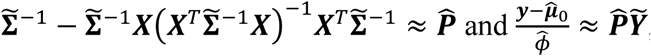, then

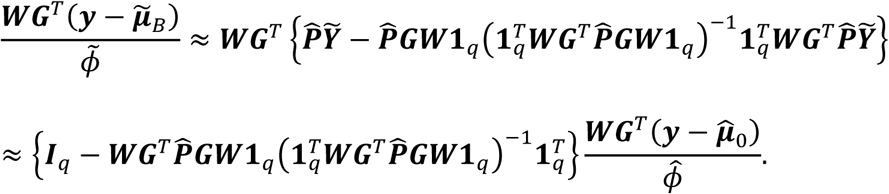

Therefore,

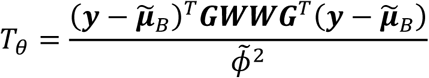

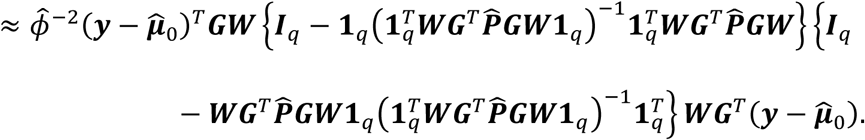

## Supplemental Data

Supplemental Data include the full authorship list with affiliations of the Trans-Omics for Precision Medicine (TOPMed) Consortium.

## Declaration of Interests

The authors declare no competing interests.

## ACKNOWLEDGMENTS

This work was supported by National Institutes of Health grants R00 HL130593 (to H.C.), U01 HL120393 (to H.C. and J.E.H.), and R35 CA197449, P01-CA134294, U01-HG009088, U19-CA203654, and R01-HL113338 (to X.L.). The authors acknowledge the Texas Advanced Computing Center (TACC, http://www.tacc.utexas.edu) at The University of Texas at Austin for providing High Performance Computing (HPC) resources that have contributed to the research results reported within this paper. Whole genome sequence analysis of fibrinogen levels in TOPMed was performed in the Analysis Commons on DNAnexus, a hosting platform that uses Amazon Web Services (AWS) to provide a cloud data management and computing environment for large genomic data projects. Phenotype harmonization and aggregation of the fibrinogen levels across TOPMed studies were supported in part by R01 HL139553. The views expressed in this manuscript are those of the authors and do not necessarily represent the views of the National Heart, Lung, and Blood Institute, the National Institutes of Health or the U.S. Department of Health and Human Services.

## TOPMed

Whole genome sequencing (WGS) for the Trans-Omics in Precision Medicine (TOPMed) program was supported by the National Heart, Lung and Blood Institute (NHLBI). WGS for “NHLBI TOPMed: Genetics of Cardiometabolic Health in the Amish” (phs000956.v1.p1) was performed at the Broad Institute of MIT and Harvard (3R01HL121007-01S1). WGS for “NHLBI TOPMed: The Cleveland Family Study” (phs000954.v1.p1) was performed at the University of Washington Northwest Genomics Center (3R01HL098433-05S1). WGS for “NHLBI TOPMed: Genetic Epidemiology of COPD (COPDGene)” (phs000951.v1.p1) was performed at the University of Washington Northwest Genomics Center (3R01HL089856-08S1). WGS for “NHLBI TOPMed: The Framingham Heart Study” (phs000974.v1.p1) was performed at the Broad Institute of MIT and Harvard (HHSN268201500014C). WGS for “NHLBI TOPMed: The Jackson Heart Study” (phs000964.v1.p1) was performed at the University of Washington Northwest Genomics Center (HHSN268201100037C). WGS for “NHLBI TOPMed: San Antonio Family Study” (phs001215.v1.p1) was performed at Illumina Genomic Service (R01 HL113323). WGS for “NHLBI TOPMed: Atherosclerosis Risk in Communities” (phs001211.v1.p1) was performed at the Baylor College of Medicine Human Genome Sequencing Center (HHSN268201500015C and 3U54HG003273-12S2) and the Broad Institute of MIT and Harvard (3R01HL092577-06S1). WGS for “NHLBI TOPMed: Genetic Studies of Atherosclerosis Risk (GeneSTAR)” (phs001218.v1.p1) were performed at Illumina Inc., Macrogen Corp., and the Broad Institute of MIT and Harvard (3R01HL121007-01S1, HHSN268201500014C, 3R01HL092577-06S1). WGS for “NHLBI TOPMed: Genetic Epidemiology Network of Arteriopathy” (phs001345.v1.p1) was performed at the University of Washington Northwest Genomics Center for the HyperGen/GENOA project (HL055673, PI: Donna Arnett/HL119443, PI: Sharon Kardia) and the Broad Institute of MIT and Harvard for the AA-CAC project (PI: Kent Taylor). WGS for “NHLBI TOPMed: Multi-Ethnic Study of Atherosclerosis (MESA)” (phs001416.v1.p1) was performed at the Broad Institute of MIT and Harvard (3U54HG003067-13S1). WGS for “NHLBI TOPMed: Women’s Health Initiative” (phs001237.v1.p1) was performed at the Broad Institute of MIT and Harvard (HHSN268201500014C). Centralized read mapping and genotype calling, along with variant quality metrics and filtering were provided by the TOPMed Informatics Research Center (3R01HL-117626-02S1). Phenotype harmonization, data management, sample-identity QC, and general study coordination, were provided by the TOPMed Data Coordinating Center (3R01HL-120393-02S1). We gratefully acknowledge the studies and participants who provided biological samples and data for TOPMed.

## Old Order Amish Study (Amish)

The Amish studies upon which these data are based were supported by NIH grants R01 AG18728, U01 HL072515, R01 HL088119, R01 HL121007, and P30 DK072488. See publication: PMID: 18440328.

## Cleveland Family Study (CFS)

Support for the Cleveland Family Study was provided by NHLBI grants R01 HL46380, R01 HL113338, and 1R35HL135818.

## Genetic Epidemiology of COPD Study (COPDGene)

This research used data generated by the COPDGene study, which was supported by NIH grants U01 HL089856 and U01 HL089897 from the National Heart, Lung, and Blood Institute. The COPDGene project is also supported by the COPD Foundation through contributions made to an Industry Advisory Board comprised of AstraZeneca, Boehringer Ingelheim, GlaxoSmithKline, Novartis, Pfizer, Siemens and Sunovion. A full listing of COPDGene investigators can be found at: http://www.copdgene.org/directory.

## Framingham Heart Study (FHS)

The Framingham Heart Study has been supported by contracts N01-HC-25195 and HHSN268201500001I and grant R01 HL092577. Fibrinogen measurement was supported by NIH R01-HL-48157. The Framingham Heart Study thanks the study participants and the multitude of investigators who over its 70 year history continue to contribute so much to further our knowledge of heart, lung, blood and sleep disorders and associated traits.

## Jackson Heart Study (JHS)

The Jackson Heart Study (JHS) is supported and conducted in collaboration with Jackson State University (HHSN268201300049C and HHSN268201300050C), Tougaloo College (HHSN268201300048C), and the University of Mississippi Medical Center (HHSN268201300046C and HHSN268201300047C) contracts from the National Heart, Lung, and Blood Institute (NHLBI) and the National Institute for Minority Health and Health Disparities (NIMHD). The authors also wish to thank the staffs and participants of the JHS.

## San Antonio Family Study (SAFS)

Collection of the San Antonio Family Study data was supported in part by National Institutes of Health (NIH) grants R01 HL045522, MH078143, MH078111 and MH083824; and whole genome sequencing of SAFS subjects was supported by U01 DK085524 and R01 HL113323. We are very grateful to the participants of the San Antonio Family Study for their continued involvement in our research programs.

## Atherosclerosis Risk in Communities Study (ARIC)

The Atherosclerosis Risk in Communities study has been funded in whole or in part with Federal funds from the National Heart, Lung, and Blood Institute, National Institutes of Health, Department of Health and Human Services (contract numbers HHSN268201700001I, HHSN268201700002I, HHSN268201700003I, HHSN268201700004I and HHSN268201700005I). The authors thank the staff and participants of the ARIC study for their important contributions.

## Genetic Studies of Atherosclerosis Risk (GeneSTAR)

GeneSTAR was supported by grants from the National Institutes of Health/National Heart, Lung, and Blood Institute (U01 HL72518, HL087698, HL112064) and by a grant from the National Institutes of Health/National Center for Research Resources (M01-RR000052) to the Johns Hopkins General Clinical Research Center.

## Genetic Epidemiology Network of Arteriopathy (GENOA)

Support for GENOA was also provided by the National Heart, Lung and Blood Institute (HL054457, HL054464, HL054481, and HL087660) of the National Institutes of Health. We would also like to thank the GENOA participants.

## Multi-Ethnic Study of Atherosclerosis (MESA)

MESA and the MESA SHARe project are conducted and supported by the National Heart, Lung, and Blood Institute (NHLBI) in collaboration with MESA investigators. Support for MESA is provided by contracts HHSN268201500003I, N01-HC-95159, N01-HC-95160, N01-HC-95161, N01-HC-95162, N01-HC-95163, N01-HC-95164, N01-HC-95165, N01-HC-95166, N01-HC-95167, N01-HC-95168, N01-HC-95169, UL1-TR-000040, UL1-TR-001079, UL1-TR-001420. The provision of genotyping data was supported in part by the National Center for Advancing Translational Sciences, CTSI grant UL1TR001881, and the National Institute of Diabetes and Digestive and Kidney Disease Diabetes Research Center (DRC) grant DK063491 to the Southern California Diabetes Endocrinology Research Center.

## Women’s Health Initiative (WHI)

The WHI program is funded by the National Heart, Lung, and Blood Institute, National Institutes of Health, U.S. Department of Health and Human Services through contracts HHSN268201600018C, HHSN268201600001C, HHSN268201600002C, HHSN268201600003C, and HHSN268201600004C. The authors thank the WHI investigators and staff for their dedication and the study participants for making the program possible. A full listing of WHI investigators can be found at: http://www.whi.org/researchers/Documents%20%20Write%20a%20Paper/WHI%20Investigator%20Long%20List.pdf.

## WEB RESOURCES

The URLs for data presented herein are as follows: Analysis Commons, http://analysiscommons.com/

DNAnexus, https://www.dnanexus.com/

GMMAT, https://github.com/hanchenphd/GMMAT

MMAP, https://github.com/MMAP

